# Inferring livestock movement networks from archived data to support infectious disease control in developing countries

**DOI:** 10.1101/2021.03.18.435930

**Authors:** A. Muwonge, P.R. Bessell, T. Porphyre, P. Motta, G. Rydevik, G. Devailly, N.F. Egbe, R.F. Kelly, I.G. Handel, S. Mazeri, B.M.deC. Bronsvoort

## Abstract

The use of network analysis to support livestock disease control in low middle-income countries (LMICs) has historically been hampered by the cost of generating empirical data in the absence of animal movement recording schemes. To fill this gap, methods which exploit freely available demographic and archived molecular data can be used to generate livestock networks based on gravity and phylogeographic modelling techniques, respectively. However, questions remain on the performance of these methods in capturing the topology of empirical networks. Here, we compare output from these network methodologies to a network constructed from either empirical data or randomly generated data. To facilitate this comparison, the spread of infectious diseases was simulated, it is this evaluation that demonstrates their potential utility to inform robust livestock disease control strategies.

The molecular network was the closest approximation to the empirical network, both in relation to topological and epidemic characteristics, whereas size of epidemics in the gravity network tended to be larger, better agreement across all three networks was observed when; a) total nodes infected, b) percentage infection take off were compared. These methods consistently identified the same important animal movement and trade hotspots as the empirical networks. We therefore consider this proof-of-concept that demographic data such as censuses and archived molecular data could be repurposed to inform livestock disease management in LMICs.

**Author summary:** Live animal movements in Africa represent a significant risk of transmission and spread of infectious diseases in livestock populations, and therefore, have direct implications on the food security of the continent. Here we explore the potential utility of available data to support control strategies, by comparing movement networks inferred from such data i.e. census and pathogen molecular data using gravity modelling and phylogeography respectively. Their utility is evaluated by comparing their topology and disease spread characteristics to empirical live animal movement. Based on our results, we posit that archived data can be repurposed to support infectious disease control on the African continent.

## Introduction

Network-based approaches enable users to describe livestock movements as a spatial network where nodes may represent villages, markets, herds, or individual animals, whereas edges define the movement of at least one animal from one node to another [1,2]. The utility of this approach has been exploited for epidemiological purposes, in particular, to understand the role of live animal movements in disease spread and control strategies [3–5]. While the use of network methods has predominantly been in developed countries due to presence of large live animal movement traceability data sets [6], relatively little has been done in sub-Saharan Africa (SAA) due the paucity of traceability data [7].

To date, empirical livestock movement networks in SSA have been constructed from purpose-made surveys, records of official permits granting movement licences or gathering trade records from key locations [8,9]. However, such data gathering efforts are expensive and may lead to data restricted in scope and most often of poor quality. Our previous work has re-constructed a high quality live animal network from such data in Cameroon but it took two years to gather the data and eight months of cleaning and analysis [8]. One of the approaches than can be used to infer networks is gravity modelling, here the amount of interactions that are likely to exist between any two locations are quantified as a function of their population sizes and the distance that separates them [10–13]. As such, it has been used to better understand multi-lateral dynamics behind migrations and flow of goods [10] and can be easily extended to infer a gravity network representing the movements of livestock [14] and humans [10] between populations.

Alternatively, livestock movements networks can be inferred from the interactions between livestock hosts and pathogens they carry. This approach exploits the principals of “measurably evolving populations” (MEP) of pathogens through the use of phylogeographic modelling [15]. This assumes that pathogen populations carried by livestock exhibit detectable amounts of *de novo* evolutionary changes in time and space [15]. In other words, the pathogens hitch-hiking on the hosts provides the evolutionary signal (molecular distance) and host locations provides the physical distance. A linear relationship between these two parameters determines the data points to infer a molecular livestock network. In this regard, considerable amount of molecular data has been collected over the years as part of research activities [16–18]. In particular, molecular data on *Mycobacterium bovis*, the causative agent of bovine tuberculosis which currently exist in form of spoligotypes and MIRU-VNTR types for cattle in Cameroon [16,17,19].

Leveraging such data not only advances data driven disease control but safe-guards sustainable food security on the continent. However, questions remain on the performance of methods used to capture/infer the topology of empirical networks. Therefore, in this study we aim to evaluate the performance and potential utility of these alternatives, i.e. gravity and phylogeographic modelling methods, to infer livestock movement networks and capture topological and epidemiological features that emerge from empirical networks. To do so, we used data from Cameroon as an exemplar because of its unique location and by extension role in linking cattle movements in central and west Africa[20,21].

## Materials and Methods

### Study area

Cameroon is a middle-income country in Central Africa, which strategically connects livestock in Central Africa to West Africa by trading live cattle to neighboring countries such as Nigeria, the Congo and Gabon (**Figure 1**). Cameroon is home to approximately eight million head of cattle providing nutrients to twenty-three million people in Cameroon [17]. Over three quarters of the cattle is reared in the north of Cameroon which supplies live animals for slaughter to abattoirs in the larger cities in the south as well as to some neighboring countries. Pastoral seasonal mobility and Sahel transhumant migration [20], where people and cattle move from as far east as Central African Republic through the northern half of Cameroon to as far west as Mali also occurs [20]. The study focuses on the main cattle rearing areas in the north, as this represents the majority of the cattle movements in Cameroon. Administratively, the country is divided into fifty-eight divisions and then into sub-divisions, it is the latter that form our unity of analysis.

**Figure 1.**
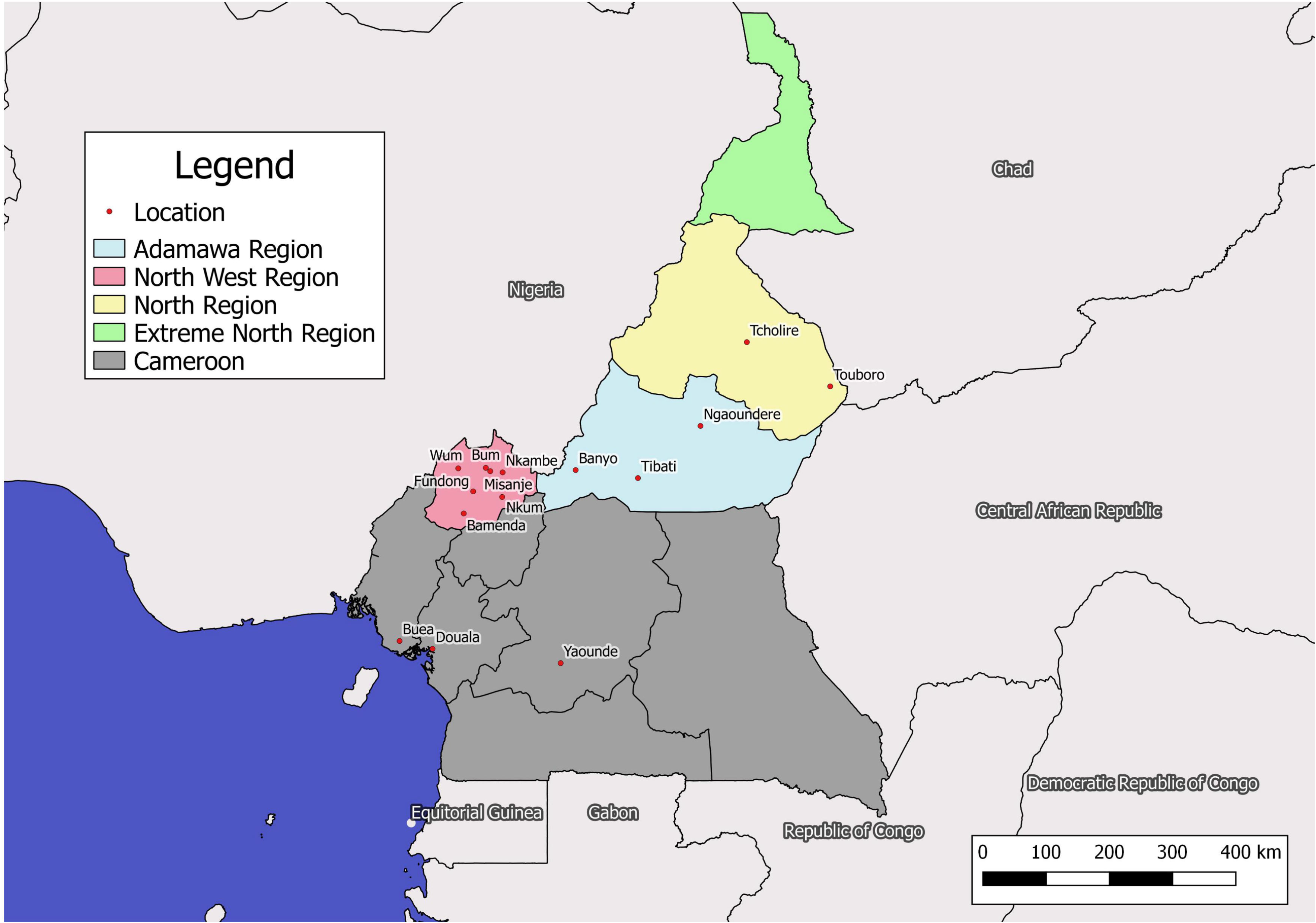
Map of Cameroon showing the cattle raring regions highlighted in blue, yellow, green and red. We also show the locations pertinent to this work. The map is plotted in Qgis version 3

### Data sources

The empirical data is available from Motta et al 2017 [7], this dataset contains 22,698, representing 9% of cattle movements in twelve months of market trading data collected in 2014-2015 from across Cameroon[7] with volumes and direction of flow between markets (**Supplementary DBSX1**). Details of the data collection procedures are described in Motta et al. [10] and in Supplementary DBSX1.

The molecular data is available from Egbe et al [17], this dataset contains spoligotyping and MIRU/VNTR data on *Mycobacterium bovis* from 2346 cattle slaughtered in four abattoirs in north of Cameroon between 2012-2014 (**Supplementary DBSX2**).

The demographic data came from the Cameroonian human and livestock census from 2005-2007 which is available from http://www.statistics-cameroon.org (**Supplementary DBSX3**).To account for the long onset of pathology in bovine tuberculosis, we allow for that a temporal lag of 4-7 years between the census and pathogen molecular datasets [22,23]. We also obtained cartographic information from GADM database of Global Administrative Areas (www.gadm.org) and merged it with respective sub-division.

### Network construction

The network construction is based on twenty sub-divisions common to all three datasets. These sub-divisions account for 5% and 20% of the human and cattle population of Cameroon (**Supplementary DBSX3**).

#### The Empirical Network

Full details of the methods for the development of the empirical network are described in Motta et al. [10]. Briefly, a weighted static directed network was built. Markets formed the nodes of the network and the links between markets were defined as the movement of at least one animal from an origin market to a destination market. For the purposes of this current analysis, data from different market were aggregated at sub-division level, i.e. network nodes were sub-division centroids [10].

#### The Phylogenetic Network

The phylogenetic network was constructed using molecular data for *M. bovis*, a slow evolving pathogen that is endemic in Cameroon [17]. *M. bovis* genotypes were defined as a pattern generated from spoligotyping and a standard 24 Loci MIRU-VNTR [24]. A mutation event was defined as a difference in a MIRU-VNTR type between any two isolates with the same spoligotype [24]. The molecular distance between any two *M. bovis* genotypes was then computed as the number loci to which they were different. The physical distance between the *M. bovis* samples was calculated using the Euclidean distances[25] between the centroids of the sub-division from which the *M. bovis* sample was obtained. Here we assumed that; a) animal-pathogen data with similar temporal and spatial scales contain detectable “historical record/signal of movement”, b) there is a linear relationship between physical distances of cattle (host-host transmission) and molecular distances (pathogen mutation events) [15]. It is this relationship that we exploit to generate the molecular network i.e. all data points that satisfy this linear relationship. The phylogenetic network is henceforth referred to as the molecular network and at this stage, the generated network is undirected, we then introduce directionality in the network using the difference in *M.bovis* genetic diversity between a pairs since any location with a high diversity of *M.bovis* is more likely to be the source of for other locations [17]. A more in-depth description can be found in supplementary materials (**Figure S3&4**).

#### The Gravity Network

Gravity modelling exploits the attraction and repulsive forces between two bodies [10,26,27]. These forces are used to define probable links (i.e. animal movements) between any two sub-divisions with a specific population density of humans and heads of cattle. The attractive and repulsive forces are modelled using a modified gravity formula (eq1); where the gravitation force (*δ*) here represents the attraction of cattle as a protein source for a given human population in a sub-division (**Figure S2&5)**.

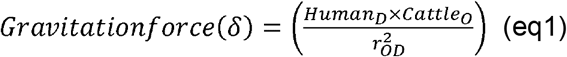

Where *Cattle_O_* is the size of the cattle population at the sub-division of origin, *Human_D_* is the size of the human population in destination sub-division and 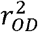 is the square distance between the sub-divisions (origin and destination).

In order to estimate the relationship between*δ* and cattle movements, we define an optimal threshold by fitting values of *δ* with presence of a link in the empirical network **Figure S6**. Since we expect an overlap in movements reflected by these two datasets, we then determine the threshold that best reflects an empirical link. This was evaluated by maximising the estimate of the area under the curve (AUC) comparing the presence of links in the network constructed using the gravity model with those present in the empirical network. To examine the practical utility of the gravity modelling approach in the absence of empirical surveillance data, we use the full census data of Cameroon (2005-2007) to generate a country wide cattle movement network. We used the optimised thresholds of *δ* from the gravity modelling framework as described in the methods section. The resultant directed network topology was then plotted on the Cameroon map using the ggmap package in R version 3.6.

#### Random network topology

To evaluate the extent to which different network inference methodologies reflect systematic differences rather than random noise, a set of random networks were generated with equivalent characteristics to that of the empirical network [28]. We use the Erdos-Renyi model for random sampling of networks and fixed the network parameters (i.e. number of edges and nodes) equivalent to the parameters of the empirical, gravity, and molecular network topologies [28]. We generated 1000 random equivalents of each network [28].

### Networks comparisons

The network-level metrics, described below, were used to compare the phylogenetic and gravity networks to the empirical network. In addition, diseases spread between nodes of these networks was simulated to gain insight on how well each network generating methods may captures “specific epidemics parameters” (Time and number of nodes affected at epidemic peak) and “non-specific epidemics parameters” (e.g. total number of nodes infected) epidemic features relevant for informing mitigation strategies.

#### Network and node level measures

The topological measures computed to describe the structure of the networks[29] included: the total number of links existing in each network, the network density *(d)*, average shortest path length *(PL)*, diameter *(di)* and the clustering coefficient (CC) These are defined in **Table S1**.

We also calculated node-level metrics for each network including; in-degree, out-degree, betweenness centrality and eigenvector centrality. These were then used to characterise the role of each node.

To define the role of each node, we characterised them as; (a) “gate-keeper”, characterised by a high betweenness centrality with a low eigenvector centrality, (b) “pulse-taker”, a well-connected node characterized by a high eigenvector centrality and low betweenness centrality, or (c) a “dual function” node, characterized by both a high eigenvector and high betweenness centrality [7].

To evaluate the degree of information provided by the variability characterized between networks, we compared (topologically, network indices and overlap analysis) the three networks’ metrics with those computed from a set of 1000 randomly generated networks with the same number of edges and nodes as the considered networks. Here, all networks were generated and analysed using the ‘igraph’ package in the statistical software R version 3.6.

#### Temporal representativeness

The steps above will have generated static molecular and gravity networks which represent all paths defined in a specific temporal window[30]. In reality all paths do not exist at all times, so static networks tend to create accessibility pathways in the network which are not always there, such an error is quantifiable as a comparison between the number of paths in a static network and a time-series network[30]. In this case, we split the empirical network into Quarters 1 to 4 of a year as reported in [7]. We then define temporal fidelity (C) as be shown below;

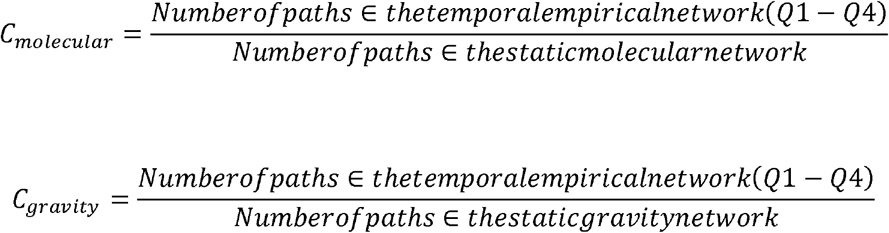

A temporal fidelity(C) of 1 means the temporal network is well represented by the static network, on the other hand a lower (C) means the opposite. The causal error defined as overall overestimation of disease outbreaks if the static network is use is approximated as 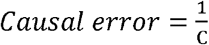

#### Disease transmission potential for networks

To evaluate the disease implications of the contact structure inferred by the generated networks, we simulate disease spread using a discrete SEIR model in *Epinet* package. The SEIR model included parameters for the infection rate (i.e. the probability of contagion after contact per unit time, β), the recovery rate, (defined as the probability of recovery per unit time, γ), and the rate of removal of infectious nodes (θ_i_).

To seed the infection, we fixed β = 0.3, γ=1 and θ_i_ as this was the lowest parameter values at which the disease propagates through the network regardless of where it is seeded on the 20 nodes. The disease spread simulation was repeated for 200 separate runs. The latency period was set to zero. In order to evaluate real-life scenario, we repeated the above simulation across a range of parameters i.e. β = 0 - 0.31 and γ = 0 - 2 simulating rapidly spreading diseases such as foot-and-mouth disease (FMD) and contagious bovine pleural pneumonia (CBPP) in a super herd setting [31,32]. In all cases, a fixed recovery rate, θ_i_ = 5, was used. The same process was done for each of the random networks. We analyse the overlap of epidemic characteristics from each network presented as a density plot in *ggplot2* package in R version 3.6.

We also generated 1000 random-equivalents for each network topology and repeated the simulation process on each network [28]. The coefficient of overlap between networks[33] on a density plot for “specific” epidemic parameters was assessed between the empirical and all the other network topologies including the random network. Total overlap was interpreted as 100% agreement between networks. Furthermore, we examined the impact of removing a node on disease spread in each of network. Here we simulate disease spread (β = 0.2, γ = 1, θ_i_ = 5) and sequentially dropping one node and seeding the infection at each of the remaining nodes for 200 iterations. We then compare epidemic statistics from these simulations with dropped nodes to a baseline full network. The epidemic statistics here were the mean number of infected nodes and the percentage of model runs where there was take off (>1 node infected)

## Results

### A comparison of network and node level measures

**Figure 2** highlights the topological similarities, in term of presence and direction of flow, of the inferred Gravity network and Molecular network with the empirical network. However, Gravity network contained more edges (routes) when compared to empirical networks and Molecular network. The gravity network (134) captures nearly three times as many edges as the empirical (43) and twice as many as the molecular networks (53), **Table 1**.

**Figure 2.**
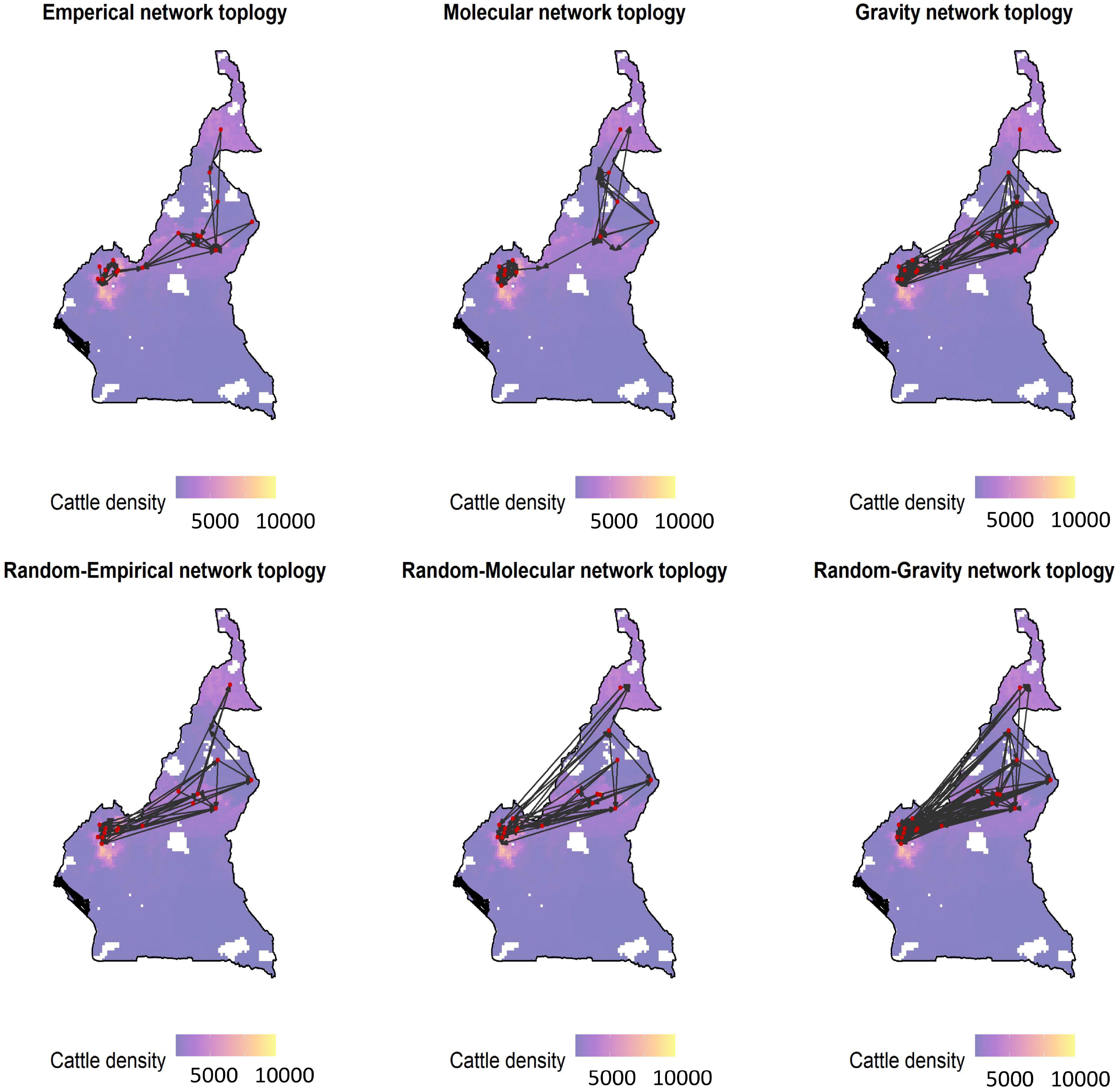
Generated networks i.e. empirical, molecular gravity and the random network plotted on a base map from Open Street Maps using *ggmaps* package in R.

**Table 1:**
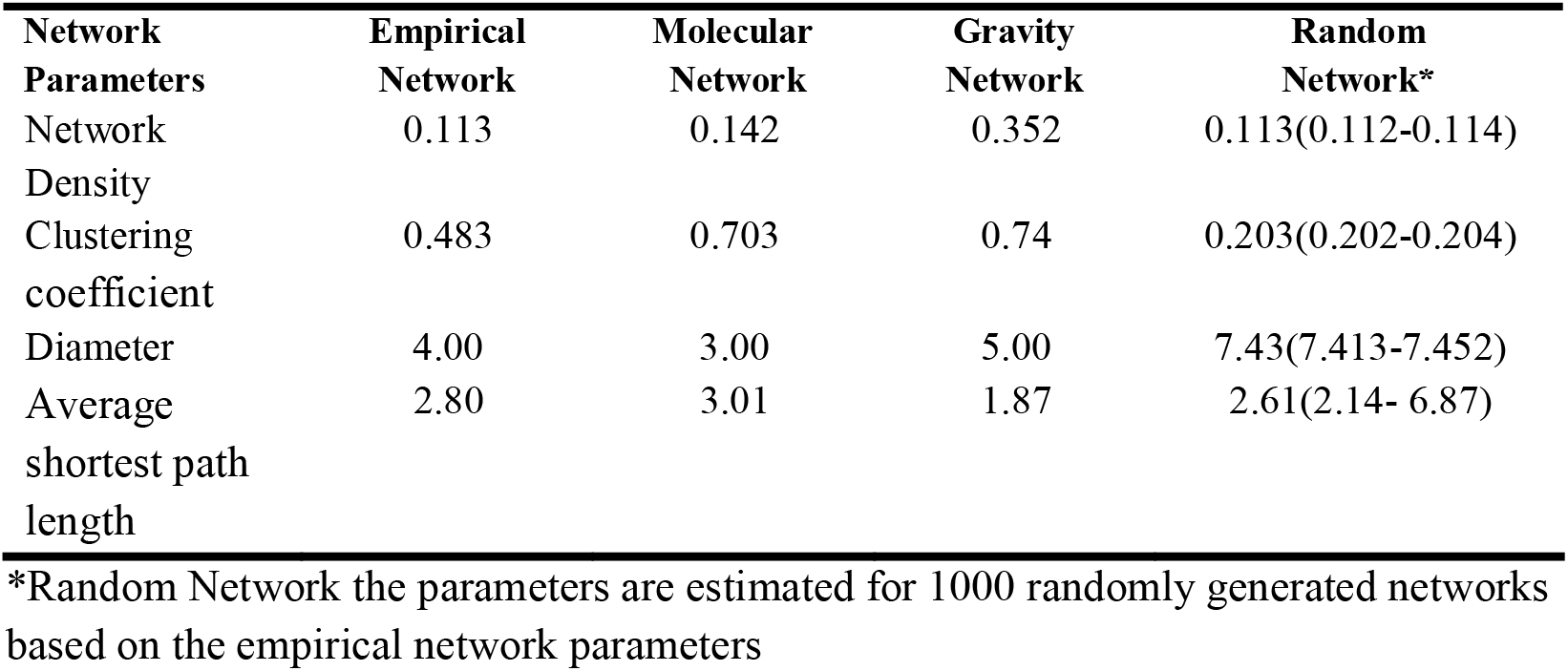
The comparison for network characteristics

A comparison of un-directed and directed networks shows that 40% and 22% of edges in molecular network are present in the empirical network, respectively. Similarly, the gravity and empirical networks share 41% and 27% of undirected and directed edges, respectively (**Table S1**). However, better agreement is observed between molecular and gravity i.e. 64.8% and 92.5% in directed and un-directed networks respectively. Interestingly, all three networks identified the same critical link between cattle populations of Cameroon. On the other hand, all three random networks exhibited topological characteristics which were considerably different from their respective network equivalents. For example; route in terrain that could not support animal movement (**Figure 2 & Table 1**).

Compared to the density of the empirical network (*d* =11%), the molecular network was of similar density (*d* =14%) whereas the gravity network was denser (*d* =35%). While these findings highlight a higher clustering coefficient (CC) for Molecular network and Gravity network, it is noteworthy that the former exhibits the shortest path length. In general, we observe considerable differences between these and their random equivalents on almost all measures **(Table 1)**.

By plotting the relationship between the local centrality measures eigenvector and betweenness, we can characterize the role of each node in the network (**Figure S6**). In doing so, we identify nodes that are “pulse-takers” (a well-connected node characterized by a high eigenvector centrality and low betweenness centrality), and those that have a dual-purpose (a node characterized by both a high eigenvector and high betweenness centrality). Here, Banyo and Ngaoundere played a dual-purpose role, with the former exhibiting this in both empirical and gravity networks. However, the same was not true for Banyo in the molecular network, this role is instead taken on by Fundong. We note that most “pulse-takers” were located in the North West region of Cameroon. At national level (applying the gravity model to the full Cameroonian census dataset), an emerging trend of animal movements are from Northwest and the Adamawa towards the populous cities in the south like Yaoundé, Douala and Buea is evident. Here we note that in addition to Banyo, Tibati acts as a critical link between the Northwest and the Adamawa. Furthermore, Touboro and Ngale are likely important transboundary hubs in the West and East of Cameroon respectively (**Figure 5A & B**).

**Figure 3.**
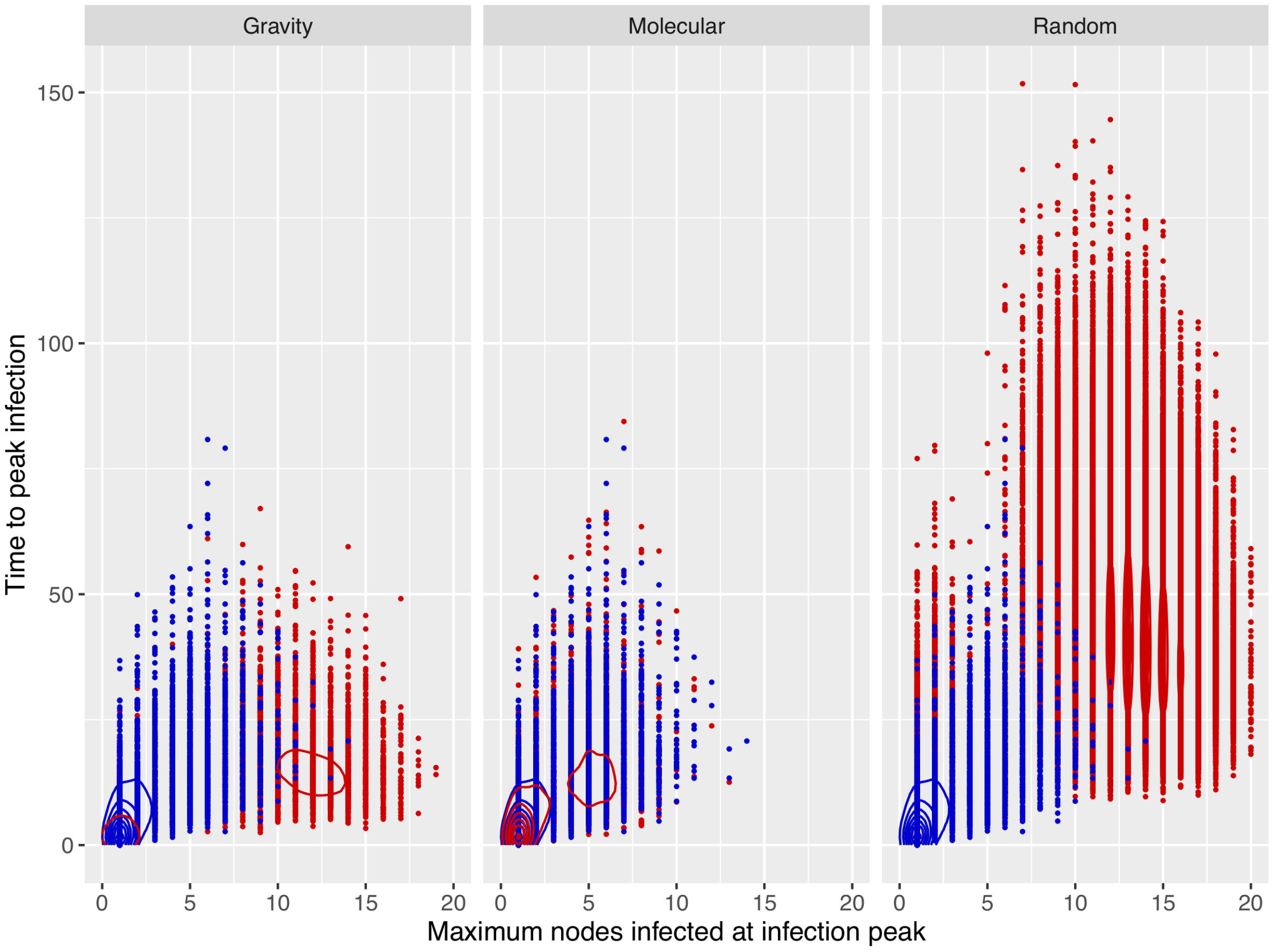
Density plot of time to peak infection and the maximum number of nodes infected. The blue and red colour represents the dynamics of the empirical and the respective network topology. This plot is generated from the output of two hundred runs per node per network topology i.e. 4000 data points per network.

**Figure 4.**
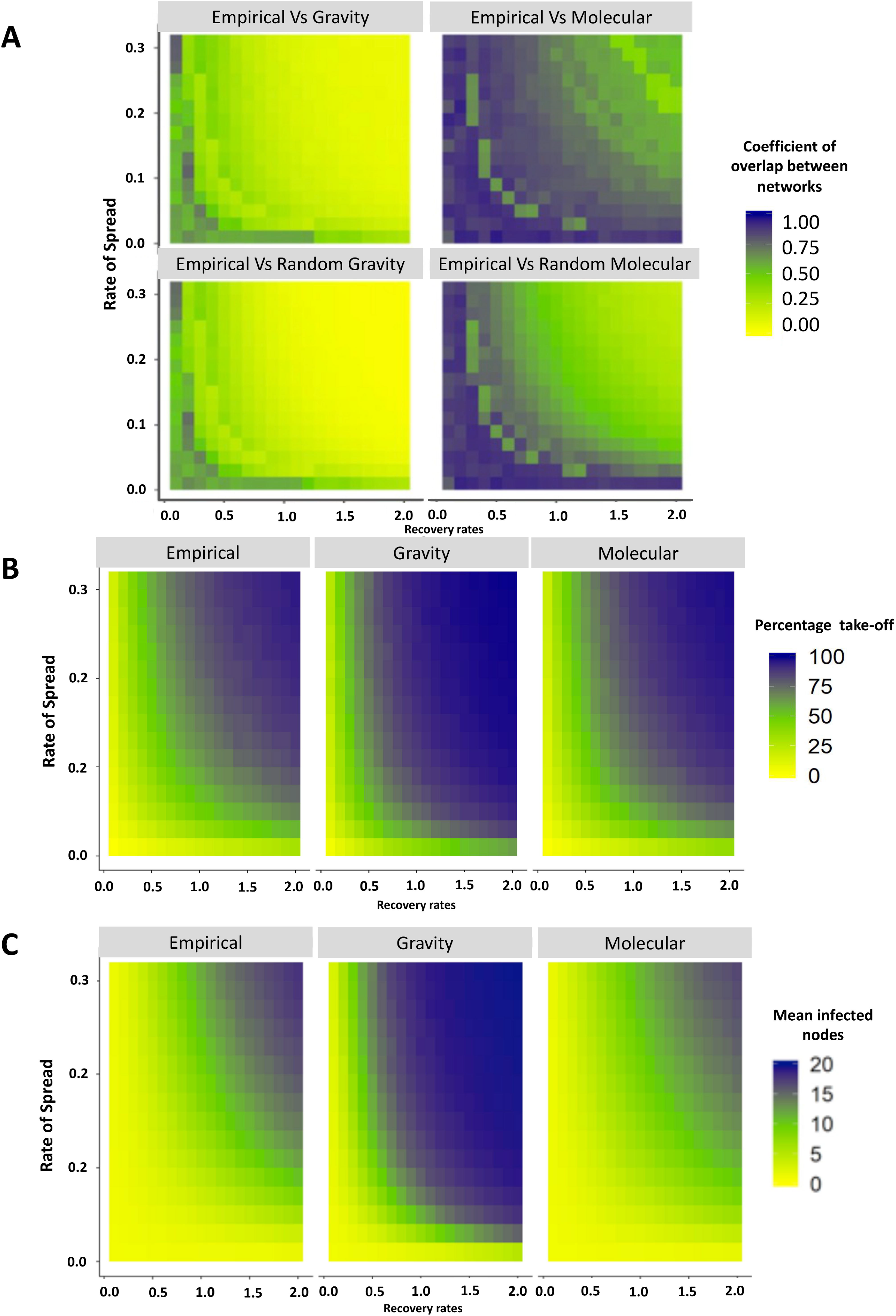
Coefficient overlap between the networks. Here we compare the empirical and its random equivalent to the molecular and empirical network dynamics. The coefficient of overlap is fitted for a range of Rate of spread (β) and γ Recovery rates (p) values on the y and x axis respectively. **B-** Coefficient overlap between the networks when we consider percentage of infections that take off and **C** is when mean number of nodes infected at the end of the infection. Here B and C represent the “non-specific epidemics parameters”

**Figure 5.**
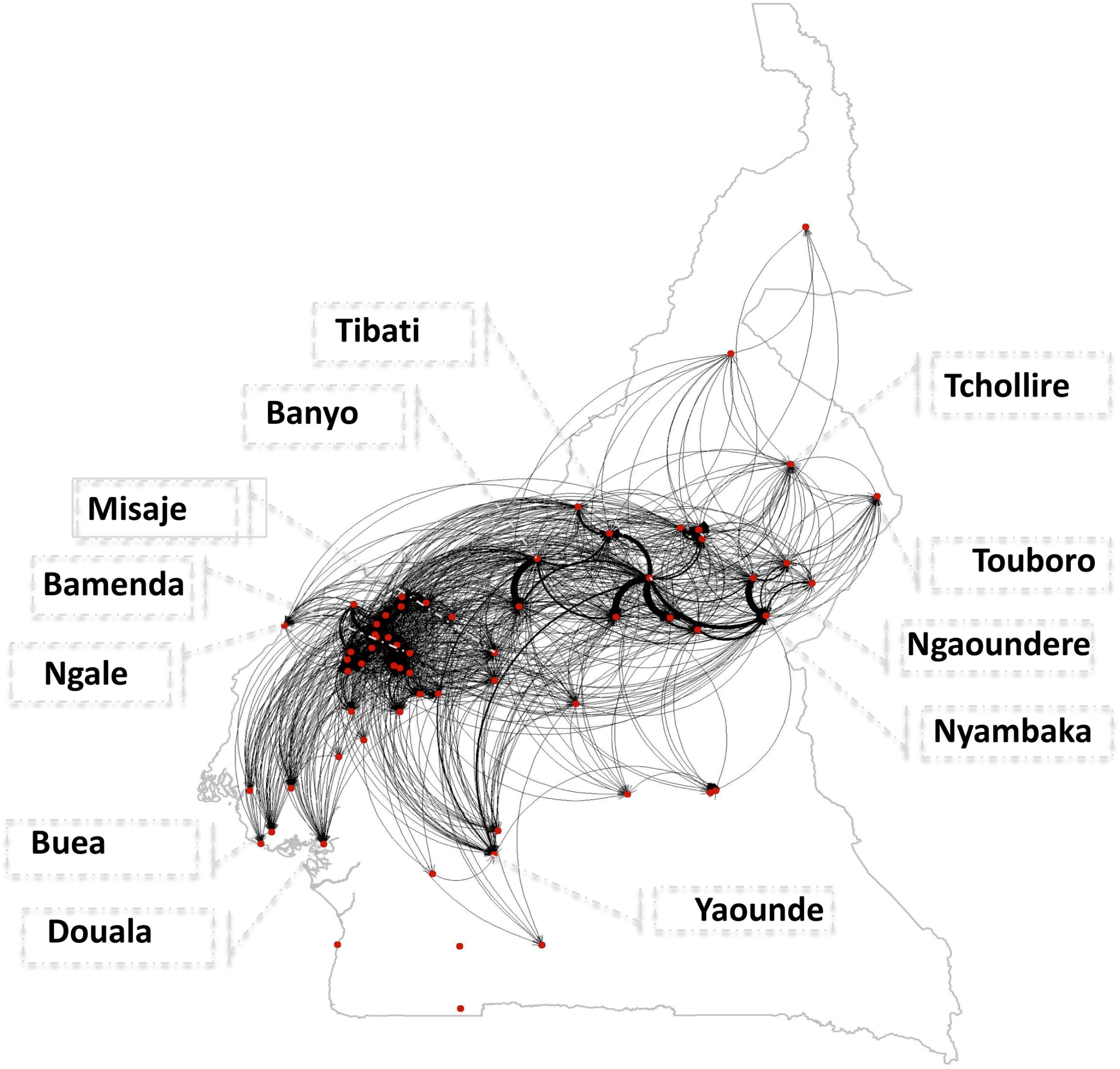
National-wide cattle movement routes generated from the demographic data using thresholds optimised for gravity model. **B**. Key actor analysis done to identify the important nodes for cattle movement

### Temporal representativeness

To evaluate the temporal representativeness of our networks i.e. if they consistently capture pathways when they exist, we compare them with a quarterly empirical network. Here we note that Gravity network’s temporal fidelity ranged between 0.201 & 0.238 while that of Molecular network varied between 0.111 & 0.222 (**Table 2**). Note that the activity of pathways varied across the year which is why the temporal fidelity, although higher than the rest, ranged between 0.76 for first quarter and 0.88 for the fourth quarter. The causal error i.e. the propensity to overestimating an outbreak when the static gravity and molecular network were used was on average at 4.5 and 5.8 respectively. This error was lowest for the static empirical network (1.17).

**Table 2:**
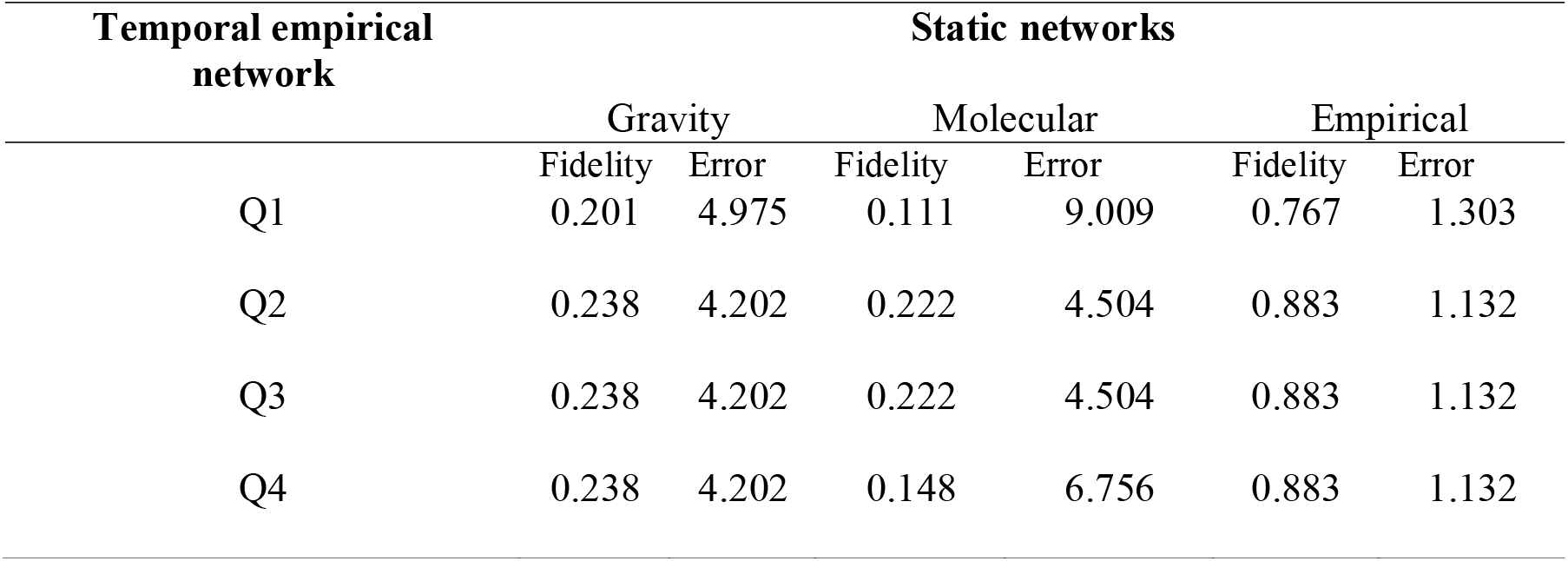
Shows the assessment of temporal fidelity

### A comparison of network disease transmission potential

Comparison based on epidemic characteristics (time to peak infection versus nodes infected at peak), showed that it took the same time to infect nodes in the empirical network, molecular and gravity network, but the later had more nodes infected at the end **Figure 3.** Indeed, the level of overlap between molecular and empirical on these epidemic characteristics was on average over 98% and approximately 59% between empirical and gravity networks. The random networks over-estimates both time to infections peak and total nodes infected when compared to the empirical. When this comparison is done over a range of epidemic parameters i.e. Rate of spread versus Rate of recovery as shown in **Figure 4A**, the overlap between the molecular and empirical network remains high, on average 0.75. Note that the higher the spread and recovery rates the lower overlap coefficients between molecular and empirical networks. The comparison between gravity and empirical exhibits poor overlap. On the other hand, when we consider epidemic parameters such as the mean number of nodes infected at the end of the epidemic and the percentage disease take off (**Figure 4B & C**), the characteristics captured are comparable. Here the overlap is higher at high spread and recovery rates

## Discussion

We set out to compare methods used to infer livestock movement networks using Cameroon as an exemplar of a low middle-income setting with limited empirical data availability. We used phylogeographic and gravity modelling to repurpose freely available and archived data. We show that molecular data from tools regarded as out-dated in developed countries infers network structures and disease characteristics similar to empirical data. In comparison, although there was lesser agreement between Gravity and empirical network, valuable higher scale inferences can be obtained from it. Therefore, repurposing data with these approaches could provide and/or complement empirical data to unravel livestock disease dynamics in developing countries.

### The molecular network and disease dynamics

The molecular network output provided the best approximation to the empirical network in terms of topology and epidemic characteristics. This finding suggests added potential of using archived spoligotype and MIRU-VNTR *M. bovis* data to infer networks [19]. Furthermore, the molecular network captured more routes than the empirical network. While the empirical network is restricted to information captured at the cattle markets, the molecular network can be used to infer unobserved events through the clonal evolutionary pathway of *Mycobacterium bovis* [15,34]. Since *M. bovis* is a slow evolving clonal bacteria, mutational changes in MIRU-VNTR and spoligotyping take between 5-12 years to occur [35,36][19]. Agreement between the empirical and molecular network suggest the following; a) there is a temporal consistency in cattle movements in Cameroon and b) bovine tuberculosis must be an endemic disease in this setting. Indeed, Motta et al[7] suggests consistency in cattle movement between markets in Cameroon [7] and it has long been suggested that bovine tuberculosis is endemic in this region [37]. The high costs of generating this molecular data hampers its utility for this purpose in LMICs, for example; it costs approximately €60 to genotype a sample using spoligotype and 24 loci MIRU-VNTR [17], this unit cost would be too high for the tool’s routine use for surveillance. However, where this data exists in archives possibly generated through research or surveillance, it can be repurposed to support regional and national livestock disease control. Note that we propose its use at coarse rather than a fine scale because our method involved merging animal market data per subdivision.

### The gravity network and disease dynamics

The gravity network captured nearly three times the number of routes as the empirical network. Since the gravity network is generated from census data, it likely represents a wide range of movements across an animal’s whole life time hence the high number of generated movements/edges. However, not all the routes/edges captured by the gravity network will be actual movement routes. This false positive limitation has been reported elsewhere [12,13], this could be minimized by fitting, socio-economic and environmental parameters to improve the model predications[5,10,38]. However, this model refinement was beyond the scope of this work.

The difference in number of routes between the gravity and empirical networks is characterized by a low level of overlap of epidemic characteristics, particularly when we consider epidemic parameters such as size and speed to peak infection. This could be because we assumed that all movements in the gravity networks are driven by the demand of the population of consumers, i.e. the human demand for meat and dairy products being the central pull for cattle in the region. In reality, cattle move due to a variety of reasons including, the search for pasture, breeding, political insecurity as well as trade [20]. We however observe better agreement when epidemics parameters such as the total size of epidemic or proportion of infections that take off are considered, in such cases the gravity network captures similar characteristics. This therefore suggests that the utility of this network ought to be limited to non-specific aspects of disease control and management.

The large number of routes the gravity network generates inherently comes with some level of reduced specificity. Therefore, prudence should be exercised when interpreting such results. In particular, we note that the gravity network can fail to capture key nodes because of the alternative routes it generates around these key nodes. It is more than likely that the gravity network provides a worst-case scenario for preparedness, i.e. provides guidance on the maximum required resources to manage an outbreak. Refining these broad inferences requires extra-information, in this case a molecular or empirical network would be needed to guide specific strategies like nodes to target for disease control. It is noteworthy that the empirical network was used as a reference for optimizing the gravity model, this practically means the utility of the former requires some basic empirical data. The same is however not true for the molecular network, and its close approximation to empirical, means the molecular could ideally be used instead to optimized the gravity model in the absence of empirical data.

### Validity of inferences

Static networks such as the one generated here suffer a weakness of representing movement paths/routes that are not always present [30,39]. To evaluate the size of this error in our networks, we calculated their temporal fidelity which suggest that the first and last quarter represent unique characteristics given the lower fidelity. In this regard the overall overestimation error was 4.5 and 5.8, interestingly the gravity slightly out-performing the molecular network.

### Practical utility of molecular and gravity networks

Practically, the gravity model could be a good alternative to extract contact information in settings where surveillance level is limited or absent, and enable the development of the early response strategies against future livestock disease outbreaks. Where molecular data exists, it could complement empirical data to refine such strategies, i.e. where empirical data is incomplete the molecular network can fill in gaps based on pathogen mutation changes. Combined, these three layers of data not only highlight the potential to cost effectively improve responses and preparedness to livestock outbreaks but also provides a foundation for data driven disease management in countries like Cameroon. We note with interest that all generated networks show that animals move from the north regions to provide protein to the large cities in the south, an observation previously reported in Cameroon [7]. Furthermore, the methods show potential to identify important trade and transboundary animal movement hubs such as Ngale, Tchollire and Touboro [7,16,17]. We however observe limited movements in the northern part of Cameroon especially in Garoua, likely because the region has approximately the same human and animal population. This suggests, that in areas where human and animal populations are approximately the same size gravity network performs poorly. Crucially, the gravity model exploits livestock protein demand, therefore its network topology inherently reflects livestock economic forces [13] so, overlaying epidemic simulations on this provides a unique opportunity to combine trade forces with epidemiological disease information. Infectious diseases cost African economies billions of dollars in reduced productivity, lost revenues due to quarantine and restricted market access [31,32]. By exploiting the contact structure inferred by our generated networks, we demonstrate potential to utilise molecular and gravity modelling to inform diseases preparedness and control strategies[6]. This is of particular value especially because the molecular and census data used here is considered obsolete and archived respectively in developing countries.

We recognise the potential bias that could arise as result of using a network work with fewer nodes, this was on balance done to ensure the same node were compared across all networks. However this bias mitigated by the by comparison with random equivalent networks [7,29].

## Conclusion

To conclude, we have adopted method which exploits freely available demographic and archived molecular data to develop livestock network which could be used for livestock disease control in developing countries. We show that the molecular network which is a close approximation to the empirical network can be generated from data considered obsolete in wealthier countries. The gravity model captures the largest proportion of movements but likely overestimates movements and disease epidemics. Better agreement is achieved across all three networks if less specific epidemic characteristics such as the size of outbreak are investigated. Moreover, these networks identify the same important animal movement and trade hotspots. We therefore consider this proof-of-concept that archived census and molecular data could be repurposed to inform livestock disease management in LMICs.

## Supporting information

https://doi.org/10.7488/ds/2780

## Acknowledgements

The primary data used in this work was generated with funding form Wellcome Trust (WT094945) with MB as the Principal investigator. AM time was funded by Core institutional budget for the Rosin Institute strategic programme (BB/J004235/1; BBS/E/D/20002172) and later by his BBSRC Future leader Fellowship. We are also grateful to the staff of MENIPIA especially the veterinarians and delegates who diligently supported the primary work that generated this data.

---

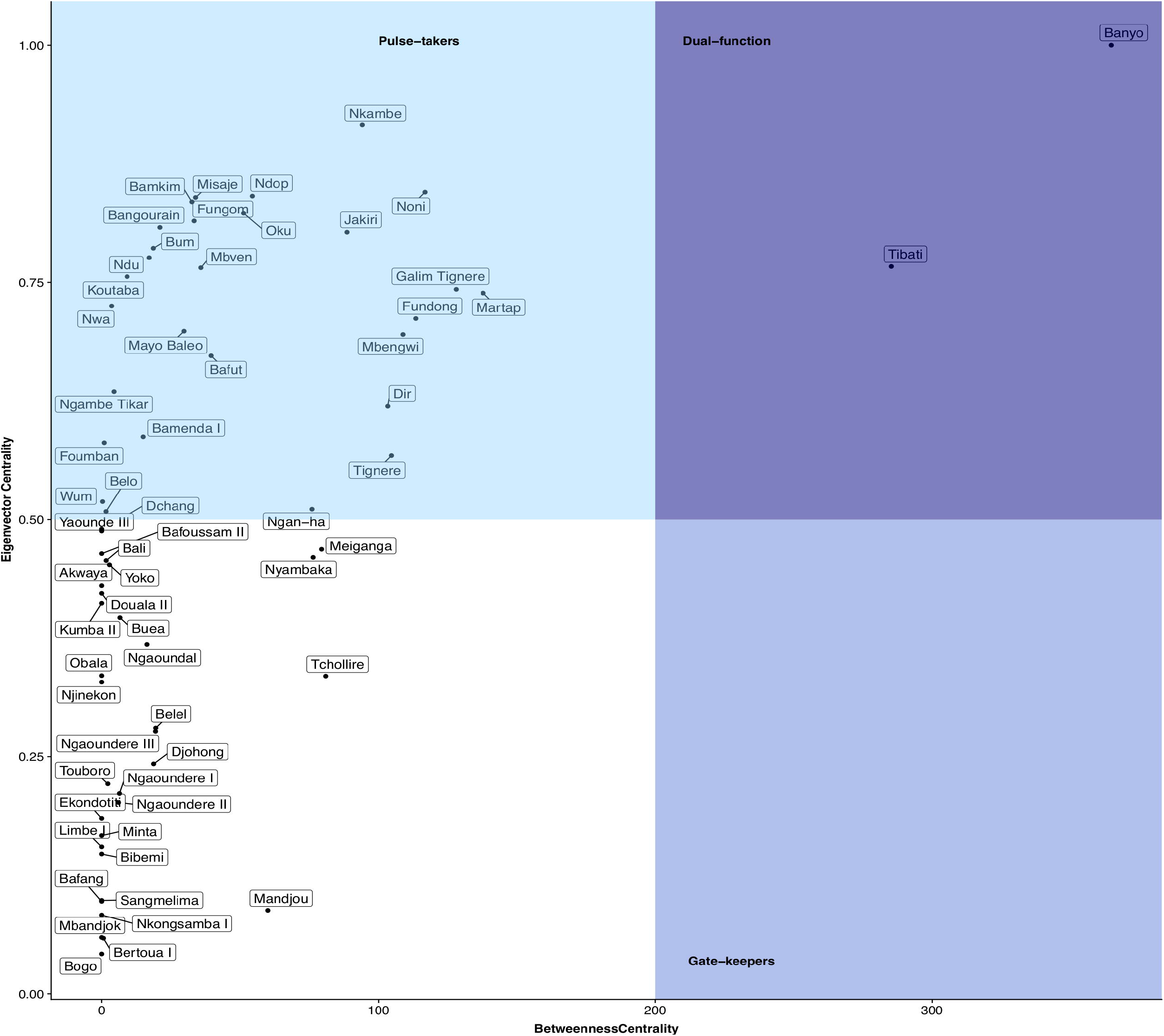

## References

1. Luke DA, Harris JK. Network Analysis in Public Health: History, Methods, and Applications. Annu Rev Public Health. 2007. doi:10.1146/annurev.publhealth.28.021406.144132

2. Efthimiou O, Debray TPA, van Valkenhoef G, Trelle S, Panayidou K, Moons KGM, et al. GetReal in network meta-analysis: a review of the methodology. Res Synth Methods. 2016. doi:10.1002/jrsm.1195

3. Moslonka-Lefebvre M, Gilligan CA, Monod H, Belloc C, Ezanno P, Filipe JAN, et al. Market analyses of livestock trade networks to inform the prevention of joint economic and epidemiological risks. J R Soc Interface. 2016. doi:10.1098/rsif.2015.1099

4. Green DM, Kiss IZ, Mitchell AP, Kao RR. Estimates for local and movement-based transmission of bovine tuberculosis in British cattle. Proc Biol Sci. 2008;275: 1001–5. Available: http://www.pubmedcentral.nih.gov/articlerender.fcgi?artid=2366193&tool=pmcentrez&rendertype=abstract

5. Chaters GL, Johnson PCD, Cleaveland S, Crispell J, De Glanville WA, Doherty T, et al. Analysing livestock network data for infectious disease control: An argument for routine data collection in emerging economies. Philos Trans R Soc B Biol Sci. 2019;374. doi:10.1098/rstb.2018.0264

6. Schrödle B, Held L, Rue H. Assessing the Impact of a Movement Network on the Spatiotemporal Spread of Infectious Diseases. Biometrics. 2012. doi:10.1111/j.1541-0420.2011.01717.x

7. Motta P, Porphyre T, Handel I, Hamman SM, Ngu Ngwa V, Tanya V, et al. Implications of the cattle trade network in Cameroon for regional disease prevention and control. Sci Rep. 2017. doi:10.1038/srep43932

8. Paolo M, Ian H, Saidou M H, Rob C, Victor N, Vincent T, et al. Implications of the cattle trade network in Cameroon for regional disease prevention and control. Nat Sci Reports. 2016.

9. Dean AS, Fournié G, Kulo AE, Boukaya GA, Schelling E, Bonfoh B. Potential Risk of Regional Disease Spread in West Africa through Cross-Border Cattle Trade. PLoS One. 2013. doi:10.1371/journal.pone.0075570

10. Eichengreen B, Irwin D a. The Role of History in Bilateral Trade Flows. The Regionalization of the World Economy. 1998. doi:10.3386/w5565

11. Zipf GK. The P 1 P 2 D Hypothesis: On the Intercity Movement of Persons. Am Sociol Rev. 1946. doi:10.2307/2087063

12. Apolloni A, Nicolas G, Coste C, El Mamy AB, Yahya B, El Arbi AS, et al. Towards the description of livestock mobility in Sahelian Africa: Some results from a survey in Mauritania. PLoS ONE. 2018. doi:10.1371/journal.pone.0191565

13. Nicolas G, Apolloni A, Coste C, Wint GRW, Lancelot R, Gilbert M. Predictive gravity models of livestock mobility in Mauritania: The effects of supply, demand and cultural factors. PLoS One. 2018. doi:10.1371/journal.pone.0199547

14. Simini F, González MC, Maritan A, Barabási AL. A universal model for mobility and migration patterns. Nature. 2012. doi:10.1038/nature10856

15. Biek R, Pybus OG, Lloyd-Smith JO, Didelot X. Measurably evolving pathogens in the genomic era. Trends in Ecology and Evolution. 2015. doi:10.1016/j.tree.2015.03.009

16. Egbe NF, Muwonge A, Ndip L, Kelly RF, Sander M, Tanya V, et al. Abattoir-based estimates of mycobacterial infections in Cameroon. Sci Rep. 2016;6. doi:10.1038/srep24320

17. Egbe NF, Muwonge A, Ndip L, Kelly RF, Sander M, Tanya V, et al. Molecular epidemiology of Mycobacterium bovis in Cameroon. Sci Rep. 2017;7. doi:10.1038/s41598-017-04230-6

18. Ghebremariam MK, Hlokwe T, Rutten VPMG, Allepuz A, Cadmus S, Muwonge A, et al. Genetic profiling of Mycobacterium bovis strains from slaughtered cattle in Eritrea. PLoS Negl Trop Dis. 2018;12. doi:10.1371/journal.pntd.0006406

19. Muwonge A, Egbe F, Bronsvoort M, Areda DB, Hlokwe T, Michel A. Molecular Epidemiology of Mycobacterium bovis in Africa. In: Dibaba AB, Kriek NPJ, Thoen CO, editors. Tuberculosis in Animals: An African Perspective. Cham: Springer International Publishing; 2019. pp. 127–169. doi:10.1007/978-3-030-18690-6_8

20. SWAC. Promoting and Supporting Change in Transhumant Pastoralism in the Sahel and West Africa. Paris; 2007. Available: http://www.oecd.org/swac/publications/38402714.pdf

21. Cour JM. The Sahel in West Africa: Countries in transition to a full market economy. Glob Environ Chang. 2001;11: 31–47.

22. Cassidy JP. The pathogenesis and pathology of bovine tuberculosis with insights from studies of tuberculosis in humans and laboratory animal models. Veterinary Microbiology. 2006. pp. 151–161.

23. Neill SD, Pollock JM, Bryson DB, Hanna J. Pathogenesis of Mycobacterium bovis infection in cattle. Vet Microbiol. 1994;40: 41–52. Available: http://www.ncbi.nlm.nih.gov/htbin-post/Entrez/query?db=m&form=6&dopt=r&uid=0008073627

24. Reyes JF, Tanaka MM. Mutation rates of spoligotypes and variable numbers of tandem repeat loci in Mycobacterium tuberculosis. Infect Genet Evol. 2010;10: 1046–51. doi:10.1016/j.meegid.2010.06.016

25. Hua H, Xie H, Tanin E. Is euclidean distance really that bad with road networks? IWCTS 2018 - Proceedings of the 11th ACM SIGSPATIAL International Workshop on Computational Transportation Science. 2018. doi:10.1145/3283207.3283215

26. Luoma M, Palomäki M. A New Theoretical Gravity Model and Its Application to a Case with Drastically Changing Mass. Geogr Anal. 1983;15: 14–27. doi:10.1111/j.1538-4632.1983.tb00756.x

27. Bossenbroek JM, Kraft CE, Nekola JC. Prediction of long-distance dispersal using gravity models: Zebra mussel invasion of inland lakes. Ecol Appl. 2001;11: 1778–1788.

28. Barabási A-L, Albert R. Emergence of Scaling in Random Networks. Science (80-). 1999;286: 509–512.

29. Wasserman S, Faust K. Centrality and Prestige. Social Network Analysis. 2012. doi:10.1017/cbo9780511815478.006

30. Lentz HHK, Koher A, Hövel P, Gethmann J, Sauter-Louis C, Selhorst T, et al. Disease spread through animal movements: A static and temporal network analysis of pig trade in Germany. PLoS One. 2016;11. doi:10.1371/journal.pone.0155196

31. Kobayashi M, Carpenter TE, Dickey BF, Howitt RE. A dynamic, optimal disease control model for foot-and-mouth-disease:. II. Model results and policy implications. Prev Vet Med. 2007;79: 274–286.

32. Lesnoff M, Laval G, Bonnet P, Chalvet-Monfray K, Lancelot R, Thiaucourt F. A mathematical model of the effects of chronic carriers on the within-herd spread of contagious bovine pleuropneumonia in an African mixed crop-livestock system. Prev Vet Med. 2004;62: 101–117.

33. Charaudeau S, Pakdaman K, Boëlle PY. Commuter mobility and the spread of infectious diseases: Application to influenza in France. PLoS One. 2014. doi:10.1371/journal.pone.0083002

34. Tanaka MM, Francis AR. Detecting emerging strains of tuberculosis by using spoligotypes. Proc Natl Acad Sci U S A. 2006;103: 15266–71. Available: http://www.pnas.org/content/103/41/15266.full

35. Tanaka MM, Phong R, Francis AR. An evaluation of indices for quantifying tuberculosis transmission using genotypes of pathogen isolates. BMC Infect Dis. 2006;6: 92. Available: http://www.ncbi.nlm.nih.gov/pubmed/16756684

36. Reyes JF, Chan CHS, Tanaka MM. Impact of homoplasy on variable numbers of tandem repeats and spoligotypes in Mycobacterium tuberculosis. Infect Genet Evol. 2012;12: 811–818.

37. Awah-Ndukum J, Kudi a C, Bah GS, Bradley G, Tebug SF, Dickmu PL, et al. Bovine tuberculosis in cattle in the highlands of cameroon: seroprevalence estimates and rates of tuberculin skin test reactors at modified cut-offs. Vet Med Int. 2012;2012: 798502. Available: http://www.pubmedcentral.nih.gov/articlerender.fcgi?artid=3329900&tool=pmcentrez&rendertype=abstract

38. Reuben J, A. DB, M. OA. Determinants of live animals and animal products trade within the ECOWAS sub-region: A gravity model approach. J Dev Agric Econ. 2014. doi:10.5897/JDAE2013.0485

39. Lentz HHK, Selhorst T, Sokolov IM. Spread of infectious diseases in directed and modular metapopulation networks. Phys Rev E - Stat Nonlinear, Soft Matter Phys. 2012;85. doi:10.1103/PhysRevE.85.066111

