## Supplementary material for "Inferring livestock movement networks from archived data to support infectious disease control in developing countries": https://doi.org/10.7488/ds/2780

### Materials and methods

#### Hypothesis and conceptual framework

This comparative analysis was implemented with some assumption about the livestock production of Cameroon a) The cattle lifecycle includes three stages highlighted here as "Rearing on farm", "movement for Trading", ending up "at the slaughter house", b) the datasets census, empirical and molecular used here represent the three stages respectively. We then generate networks from each dataset using the following methodologies;

1. Cattle movement network derived as a function of human protein demand using gravity modelling
2. Derive an empirical network topology from an edge list generated from cross- sectional study out team conducted (**DBSX1**) and
3. Using phylodynamic modelling of host-to-host pathogen transmission network, since pathogens are considered "hitchhikers" on hosts
4. Derive a random network topology. The Empirical and random are used as controls, i.e. the former as the reference/gold standard and the latter as the negative control/null

The vast majority of network structures/topologies are a product of dynamic processes [1,2], therefore one can think of the resultant network topology as a relic of the contact structure. So, based on this contact structure we can elucidate disease spread by simulation. The novelty here is the ability to repurpose generally archived data census and molecular data. Similarity and dissimilarity in topology and simulated disease characteristics between gravity, molecular and empirical network as well as the random equivalents allow us to examine the following;

a) The amount of overlap in information captured,
b) the complementally utility from i.e. the extra information each captures,
c) how specific and non-specific each network can be. All this information can be exploited to support data driven livestock disease management especially resource allocation.

#### Description of data source and context

*Empirical data set (For R code see section-A2 in Network_Generation_Code)*

The empirical network (EN) was generated using data collated on cattle movements through the livestock trading system across Adamawa, West and North-West regions of Cameroon. The lists of cattle markets present within these regions were obtained from the Ministry of

Livestock, Fisheries and Animal Industries (MINEPIA). Combining this information with the analysis of commercial connections of between markets in each region identified a total of 59 livestock markets [18].

*Census data summary*

The census data used represents approximately 8.85 and 10.3 million head of cattle and humans respectively. The human and cattle population difference between sub divisions within regions is shown in **Fig S1**. It is however noteworthy that ratio of human to cattle is highest and lowest in Adamawa and central regions respectively. Furthermore, that areas without cattle or human, or missingness of one population were excluded for our analysis.

**A** **B**

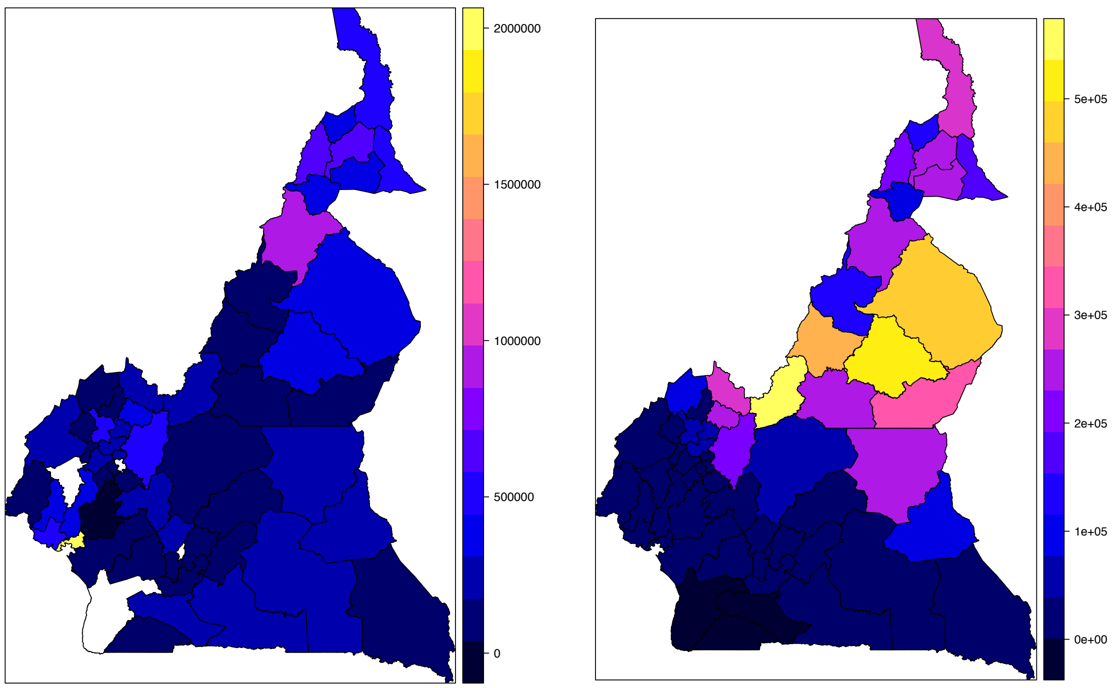


**Figure S1** Spatial distribution of the (A) human and (B) cattle population generated using ggplot in R using the Cameroonian census data of 2005-2007(**DBSX3**). Color scheme ranges from dark blue-yellow for the legend and represents population ranges (A) 0-2million and (B) 0-0.5million.

#### Principles behind each network topology

*Molecular network topology (For R code see section-A1 in Network_Generation_Code)*

Here we exploit the principals of “measurably evolving populations” (MEP) of pathogens [3] to reconstruct transmission network based of M.bovis (**Fig 2)**

**Panel 2a**

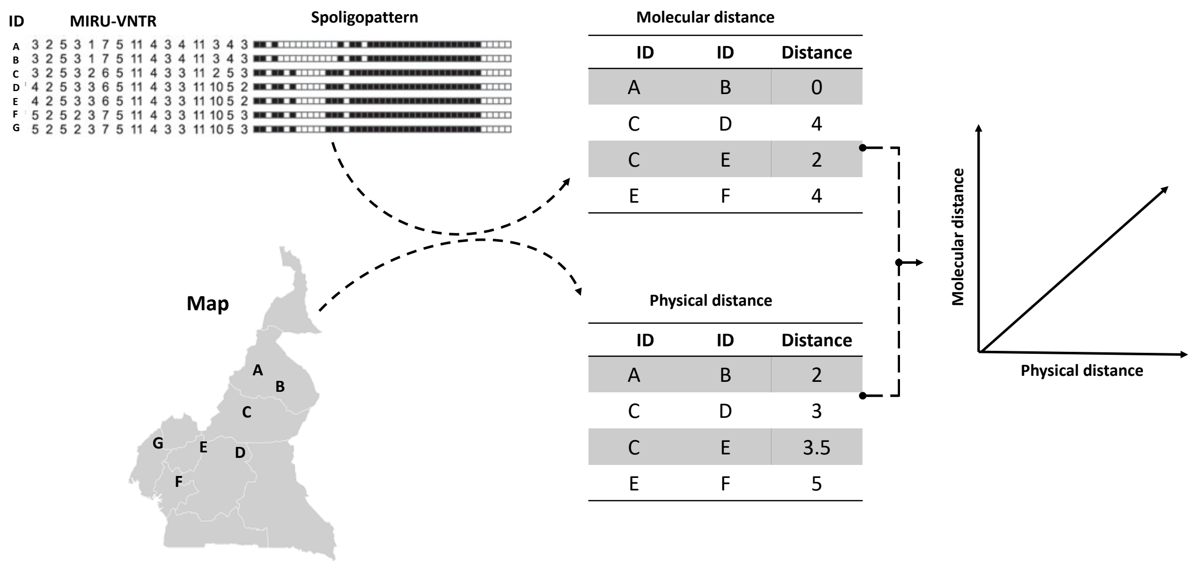


**Panel 2b** **Panel 2c**

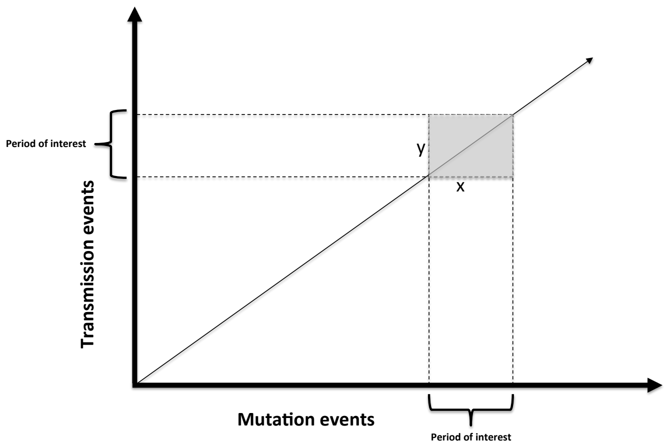


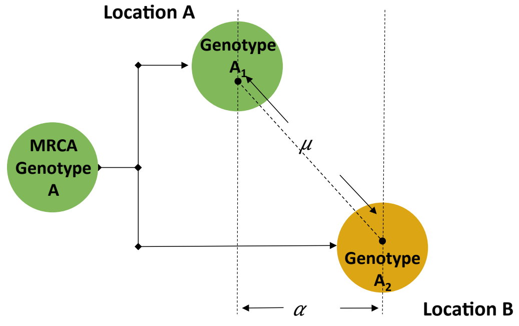


**Figure S2** illustrates the approach used to generate the molecular network. Panel A shows how we have extracted molecular distance from M.bovis genotypes and Physical distance from the host(cattle). Panel B shows how molecular distance and physical distance are related to transmission and mutation events (MEP). Here transmission events (are analogous to physical distance μ) and mutation events (analogous to genetic distance α). The "window" x-y defines the epidemiological "window" of interest in space and time [15]. Panel C puts this in phylogeographic context, the pathogen genotypes are cast in space(μ) and time(α). The two genotypes $A_1$ and $A_2$ evolved from the MRCA, most recent common ancestor, note here that they are isolated in two different locations i.e. location A & B. From phylogeography, there must be a direct relation between physical distance and molecular distance. It is the linear relationship that defines the data used to construct our molecular network

To contextualize this, take two genotypes; $A_1$ and $A_2$ recovered from cattle in location X and Y, and assuming the genotypes have a common recent ancestor $A_o$ (**Figure S2-Panel B**). We can use the molecular and physical distance α and μ to extract data points from **DBSX2** that satisfy the linear relationship (**Fig S3**) and ideally our "window" μ ~ α of interest (**Figure S2-Panel C**) & quadrant B (**Fig S3)**. Given our datasets we assume this window represents 2007 to 2014 and accounts for the period between pathogen transmission, latent infection and infectious period for the cattle in Africa. [4]. This is why we use the census data for 2005-2007. In our data set we define mutation changes as reported by [5] i.e. a mutation event as the difference in steps in a MIRU-VNTR type between any two isolates with the same spoligotype (**Fig S2-Panel A**) [16]. Physical distance is computed as linear Euclidean distances between any two sub counties. We therefore use the data points from quadrant B (**Fig S3)** to generate the undirected molecular network which we direct using the molecular diversity at each subdivision.

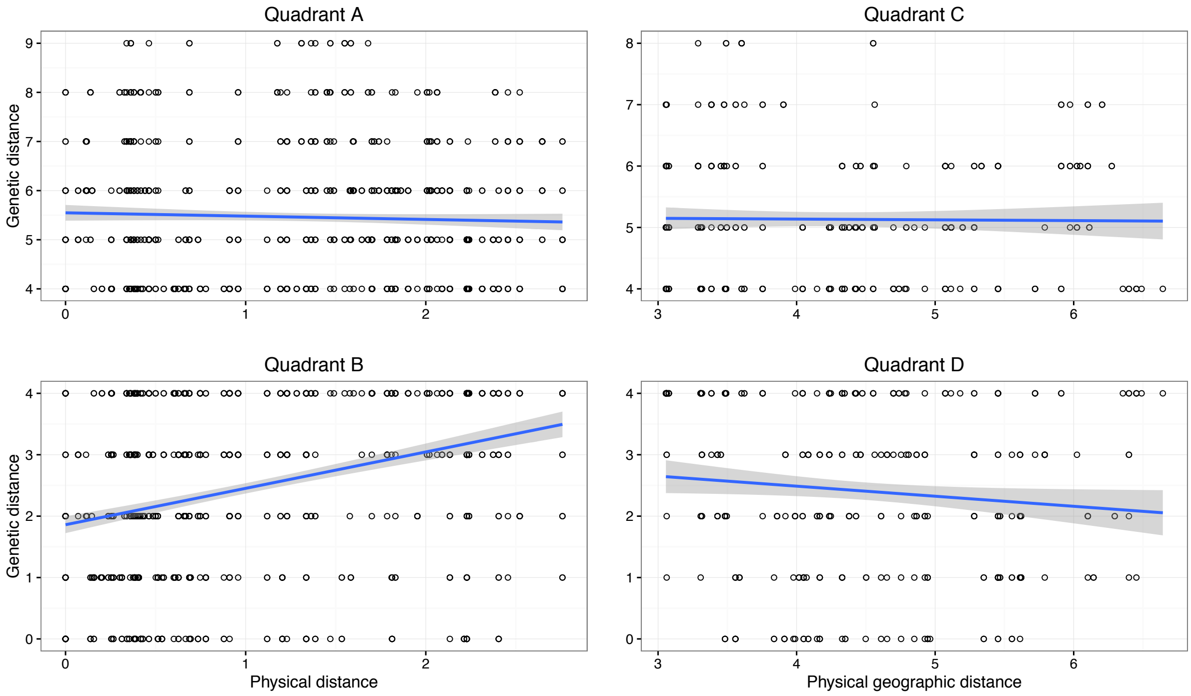


**Figure S3** Relationship between genetic distance (α) and physical distance (μ) using our data. **Quadrant B** would then represent the "window" whose data points we used to derive the molecular network topography.

The dataset used for generating the molecular distance [5] contained 25 unique subdivisions of which 20 were used for deriving the molecular network topography. Four and fifty-one unique spoligotypes and MIRU-VNTR types respectively were used to generate molecular distances. In this regard spoligotypes SB0944, SB0953, SB1025 and SB1460 listed in their order of prevalence were used.

*Gravity network topology (For R code see section-A3 in Network_Generation_Code)*

Gravity models have widely been used in economic, it is adapted from Newton's law of gravitation in physics [6], here it is modelled to measure bilateral cattle movement between any two given sub divisions of Cameroon. We assume that the movement of cattle between any two given sub division is directly proportional to their respective difference in cattle and human population and inversely proportional to the square of the Euclidian distances between them.

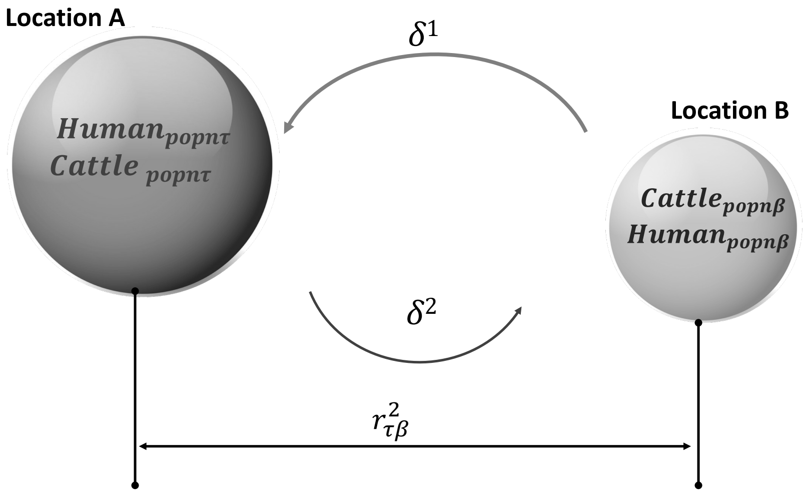


$$\delta = \left(\frac{Human_{popn\tau} \times Cattle_{popn\beta}}{r^2_{\tau\beta}}\right)$$

**Figure S4** is a gravity model illustration, the circular shapes each represents a sub division whose size represents the size of the human, cattle population. The shorter the distance between any two sub divisions, and the greater the size of the populations the greater the gravitation pull between the sub divisions hence flow of cattle between them

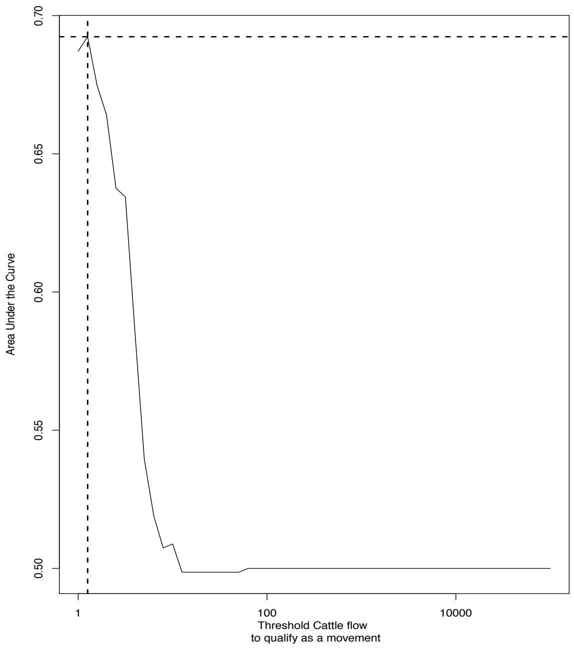


#### Gravity model threshold determination

**Figure S5** Shows the area under the curve (AUC) analysis between the gravity model and the empirical data with an aim of determining the threshold animal movement required to define an edge(link) between any two nodes in the gravity network. In this case we notice that 1.25 animals are required to define a link in the gravity model. This threshold is used to infer a national level gravity network- For R code see Cameroon_Gravity_Model.R

#### Results

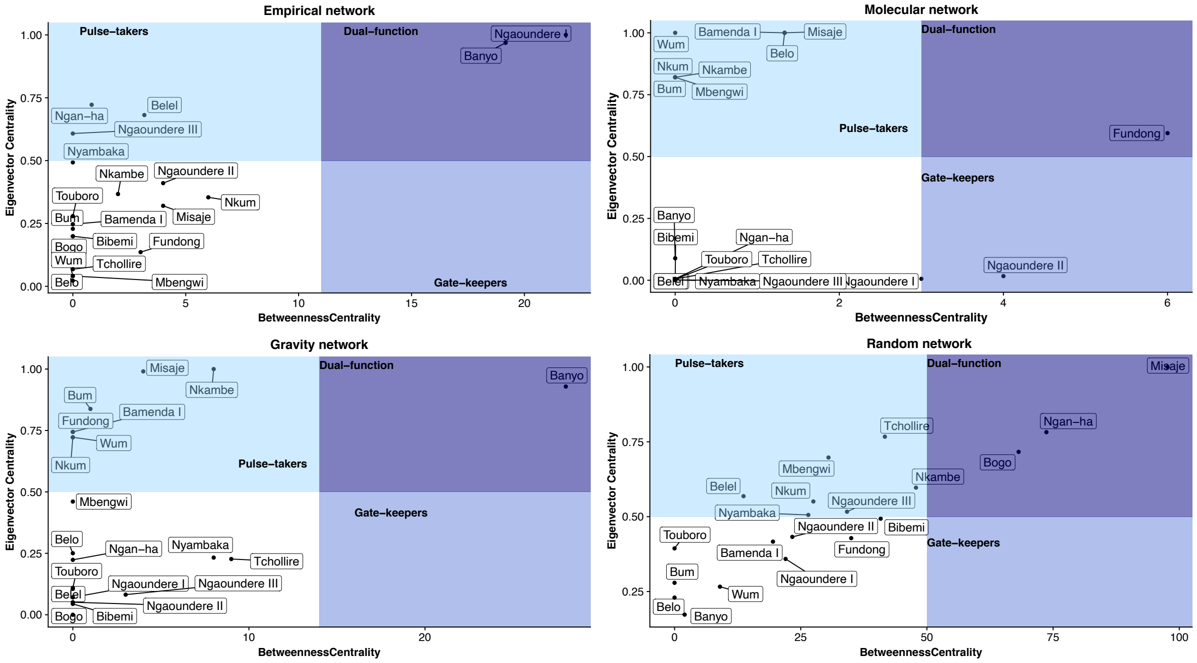


**Figure S6.** Shows the key actor analysis, this is a comparison at node level aimed at understanding the role each node plays in the networks.

As part of the comparison, we simulate infections on the generated networks as described in the materials and methods. For further details on how to implement this in R, see code Epidemic simulation_Code.R

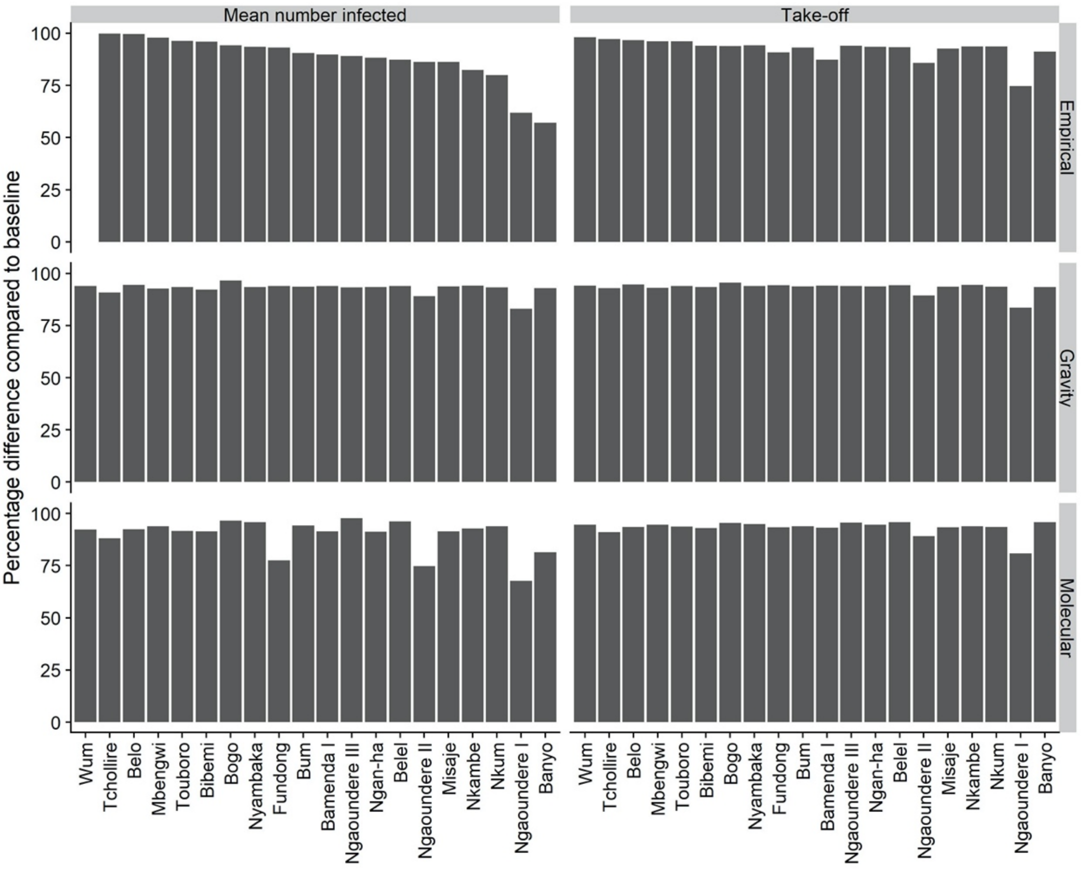


**Figure S7.** Shows the impact of removing a node mean number infected at end of infection and proportion if infection that take off in each network. This analysis aims to identify the most important node in the network *(For R code see Epidemic_Simulation_Leave_One_Out)*

Table S1: Comparison between network by presence of absence of edges

| Network 1 | Network 2 | Present n (%) Directed | Absent (n) Directed | Present n (%) Un-directed | Absent (n) Un-directed |
|---|---|---|---|---|---|
| Gravity | Empirical | 37(27%) | 97 | 101(41%) | 158 |
| Gravity | Molecular | 35(26.1%) | 99 | 136(50.7%) | 132 |
| Empirical | Gravity | 37(86.1%) | 6 | 82(95.3%) | 4 |
| Molecular | Gravity | 35(64.8%) | 19 | 100(92.5%) | 8 |
| Empirical | Molecular | 12(27.9%) | 31 | 50(58.1%) | 36 |
| Molecular | Empirical | 12(22.2%) | 42 | 44(40.7%) | 64 |

#### Terminology

Table S2: Definitions for terms used in network analysis

| Network characteristic | Definitions |
|---|---|
| Density | The ratio of the number of links in a network to the number of possible links |
| Clustering coefficient | is the ratio of existing links connecting a subdivision’s neighbors to each other to the maximum possible number of such links |

| | |
|---|---|
| Average path length | This is on average the number of steps it takes to move from one subdivision to another in each of the predicted networks |
| Diameter | This is a linear size of each of the networks, it is also defined as the longest of all shorted paths of each of the networks |
| Assortativity | Is a propensity of a subdivision to be linked with other sub divisions that are similar in someway |
| Reciprocity | Is quantitative measure in networks that describes the likelihood of subdivision in the directed networks being mutually linked |
| Transitivity | This is a measure of the extent to which a link between two sub divisions that are connect is transitive |
| **Node characteristics** | |
| Degree centrality | This is the number of links that are incident upon each of the subdivision in a network |
| Betweeness centrality | Is a measure of centrality that quantifies the number of times a subdivision in a network acts as a bridge along the shortest path between two other subdivisions |
| Eigenvector centrality | This is a measure of the influence of a subdivision in a network |
| Closeness centrality | This is a measure of metric distance between any pair of subdivisions in a network also known as their shortest path |

### References


1. Ortiz-Pelaez A, Pfeiffer DU, Soares-Magalhaes RJ, Guitian characterize the pattern of animal movements in the initial phases of the 2001 foot and mouth disease (FMD) epidemic in the UK. *Prev. Vet. Med.* **In Press,**.
2. Chaters GL *et al.* 2019 Analysing livestock network data for infectious disease control: An argument for routine data collection in emerging economies. *Philos. Trans. R. Soc. B Biol. Sci.* **374**. (doi:10.1098/rstb.2018.0264)
3. Biek R, Pybus OG, Lloyd-Smith JO, Didelot X. 2015 Measurably evolving pathogens in the genomic era. *Trends Ecol. Evol.* (doi:10.1016/j.tree.2015.03.009)
4. Reyes JF, Tanaka MM. 2010 Mutation rates of spoligotypes and variable numbers of tandem repeat loci in Mycobacterium tuberculosis. *Infect. Genet. Evol.* **10**, 1046–51. (doi:10.1016/j.meegid.2010.06.016)
5. Egbe NF *et al.* 2017 Molecular epidemiology of Mycobacterium bovis in Cameroon. *Sci. Rep.* **7**. (doi:10.1038/s41598-017-04230-6)
6. Eichengreen B, Irwin D a. 1998 *The Role of History in Bilateral Trade Flows*. (doi:10.3386/w5565)
